# NMR as a readout to monitor and restore the integrity of complex chemoenzymatic reactions

**DOI:** 10.1101/2022.05.14.491371

**Authors:** Kenneth A. Marincin, Yousang Hwang, Everett S. Kengmana, David J. Meyers, Dominique P. Frueh

**Affiliations:** Department of Biophysics and Biophysical Chemistry, Johns Hopkins School of Medicine, Baltimore, MD 21205, USA; Department of Pharmacology and Molecular Sciences Synthetic Core Facility, Johns Hopkins School of Medicine, Baltimore, MD 21205, USA; Department of Chemical and Biomolecular Engineering, Johns Hopkins Whiting School of Engineering, Baltimore, MD 21218, USA

**Keywords:** NMR, isotope filter, diffusion, chemoenzymatic reactions

## Abstract

The non-invasive nature of NMR offers a means to monitor biochemical reactions *in situ* at the atomic-level. We harness this advantage to monitor a complex chemoenzymatic reaction that sequentially modifies reagents and loads the product on a nonribosomal peptide synthetase carrier protein. We present a protocol including a novel pulse sequence that permits to assess both the integrity of reagents and the completion of each step in the reaction, thus alleviating otherwise time-consuming and costly approaches to debug and repeat inefficient reactions. This study highlights the importance of NMR as a tool to establish reliable and reproducible experimental conditions in biochemical studies.

**Graphical Abstract:** 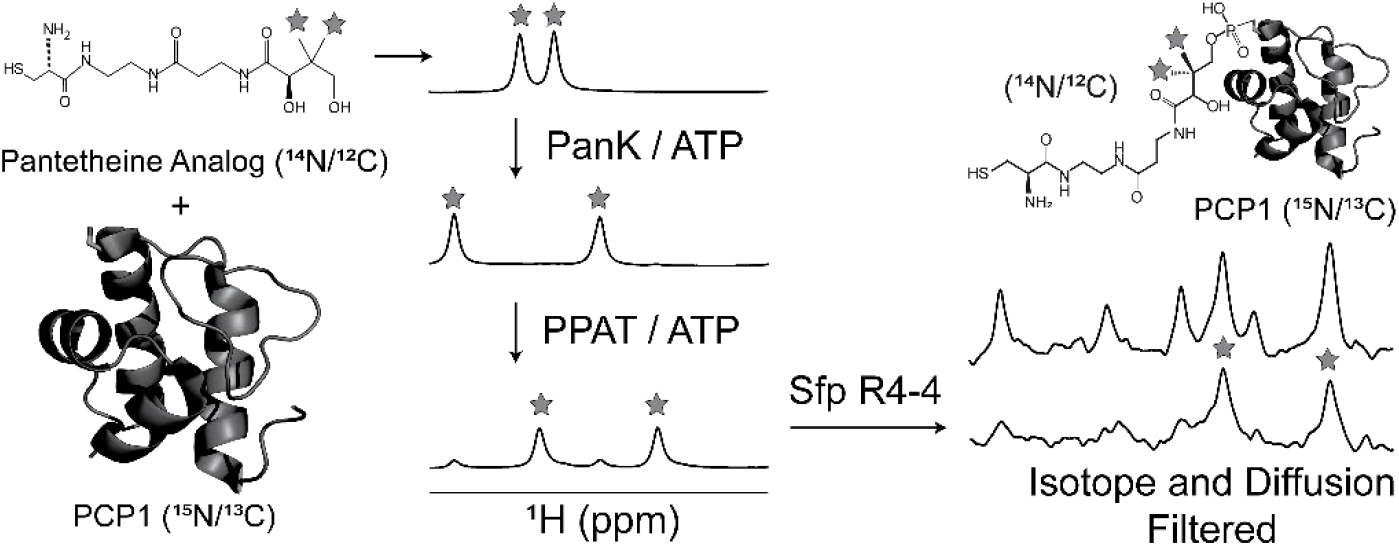

## Introduction

Nuclear magnetic resonance (NMR) promises to play a central role in determining molecular mechanisms and biological functions in near-native conditions. Recent advances in cryo-electron microscopy (cryo-EM)[1] and structure prediction algorithms such as AlphaFold[2] have greatly facilitated the determination of static structures. However, biological function and its regulation involve the timely redistribution of (transient) interactions between macromolecules, with fleeting molecular responses that are undetected by these techniques. Solution NMR provides rescue to this limitation due to its exquisite sensitivity to changes in molecular environments and its non-invasive nature. Thus, the kinetic, thermodynamic, and structural features accompanying molecular communication can be established during binding events, during biochemical reactions *in vitro*, or even in cell, resolving otherwise puzzling molecular mechanisms. Nonribosomal peptide synthetases (NRPSs) are multidomain, microbial enzymatic factories that could produce new pharmaceuticals through engineering,[3] and NMR could help explain how they operate through a dynamic architecture involving fleeting domain interactions. However, preparing samples to answer these questions requires elaborate chemoenzymatic approaches that, occasionally, lead to significant delays and may become a bottleneck to experimental studies. Here, we exploit the aforementioned advantages of NMR to identify the causes for inefficient reactions and to verify that remediations are successful.

NRPSs utilize multiple domains organized in sequential modules to covalently tether simple substrates and assemble them into a variety of complex natural products, often with medicinal (antibiotics, immunosuppressants) and industrial (surfactants, pesticides) applications.[4,5] To ensure fidelity, substrates and intermediates are attached to 20 Å phosphopantetheine (PP) moieties harbored by carrier protein domains. These domains exist in apo forms lacking PP, in PP modified holo forms, and in cargo loaded forms, with cargos including simple substrates and intermediates. Thus, synthesis occurs through a succession of interactions between carrier proteins and partner catalytic domains to introduce the PP arm, load it with a substrate through a thioester bond, and to condense this substrate to that of a CP in a downstream module through a peptide bond. In this process, intermediates are thus extended while they are transferred to downstream modules until release in the final module. This assembly-line architecture provides a basis for producing improved therapeutics by swapping and engineering domains to incorporate exogenous substrates into the final product. Unfortunately, NRPSs operate through fleeting domain and substrate interactions and, in spite of impressive crystallographic[6,7] and electron microscopy[8] studies capturing various domain interactions, the molecular determinants for substrate recognition and domain communication remain largely elusive. Structural and functional studies are further challenged by the lability of the thioester bond as substrates and intermediates often fall off during measurements. We and others have determined that PP arms and their attached cargo transiently interact with the core of CP domains,[9,10] and transient docked forms may provide a basis for dual domain and substrate recognition through encounter complexes preceding productive domain engagement in which PP arms and their cargos extend towards buried catalytic sites. One of our objectives is to determine the multiple substrate, PP, and domain binding sites in the many encounter and productive domain complexes involved in synthesis. To this aim, we rely extensively on NMR methods that exploit site-specific isotope labeling of carrier proteins, their PP arms, and their cargos. We most often employ a chemoenzymatic route to simultaneously design our labeling scheme and produce stable derivatives, in which the thioester bond is replaced by an amide bond. Critically, it is generally impossible to separate the modified and unmodified proteins during subsequent purification protocols. Inhomogeneous samples would create unnecessary challenges in NMR experiments and taint enzymatic assays or binding studies performed with other techniques. It is thus critical to obtain a nearly fully modified protein at the end of the reaction.

Carrier protein domains can be modified with a phosphopantetheine arm harboring a substrate or intermediates through a one-pot chemoenzymatic approach.[11] In this scheme (Figure 1), a series of enzymatic steps modifies a pantetheine analog to produce a Co-Enzyme A (CoA) analog[13] harboring the cargo of interest. Pantothenate kinase (PanK) first phosphorylates the terminal hydroxyl group of pantetheine before phosphopantetheine adenylyltransferase (PPAT) activates this intermediate as an adenosine monophosphate (AMP) adduct to generate a dephospho-CoA (dpCoA) derivative. Dephospho-CoA kinase (DPCK) finalizes the conversion into a CoA derivative (not shown in Figure 1), which is accepted by a phosphopantetheine transferase,[14] most commonly Sfp, and leads to a carrier protein harboring the post-translational modification of interest at a conserved residue. Traditionally, this chemoenzymatic synthesis is executed in a one-pot protocol where the carrier protein is mixed with all selected enzymes, a pantetheine analog, and ATP at once,[10,11,15–17] what we refer to in our laboratory as an all-in-one one-pot reaction. The use of the Sfp R4-4 mutant overcomes the need for DPCK.[18] However, Cryle and Co. observed that excesses of phosphoadenylates hampered the last reaction when using Sfp R4-4 and designed a sequential one-pot protocol wherein enzymes are added successively with short incubation periods of less than 30 minutes.[12] With this protocol, alkaline phosphatase from calf intestine (CIP) can be added prior to Sfp R4-4 and loaded CPs may be obtained at near complete yields. However, regardless of whether the enzymes are added successively or at once, any deviation from the expected balance in reaction kinetics governed by enzyme concentrations and their catalytic integrity may jeopardize the outcome. Notably, PanK and PPAT co-purify with CoA[19,20] that must be removed before use, and DPCK is catalytically slow and reversible.[21,22] As mentioned, Sfp R4-4 is inhibited by large concentrations of phosphoadenylates,[12] and although addition of CIP resolves this issue, incubation must be timed adequately as we will soon discuss. The take-home message from a decade of using these protocols in our laboratory and from the literature is that the protocols work adequately once they have been optimized for a given pantetheine derivative and in absence of unexpected drawbacks such as deterioration of substrate or enzyme stock solutions. However, any human error in timing or enzyme concentration may jeopardize the outcome as enzyme kinetics dictate success. Traditionally, completion is verified via matrix-assisted deionization time-of-flight mass spectrometry (MALDI-TOF MS), and SDS-PAGE provides a limited assessment of enzyme integrity and their concentration in the reaction. Importantly, these methods provide results at the outset of the reaction, which would have to be repeated if unsuccessful. Thus, time-consuming investigations may be needed to determine why substrate loading did not go to completion. To overcome these limitations, we turned to NMR as a means to monitor both the progress and integrity of this complex chemoenzymatic reaction.

**Figure 1.**
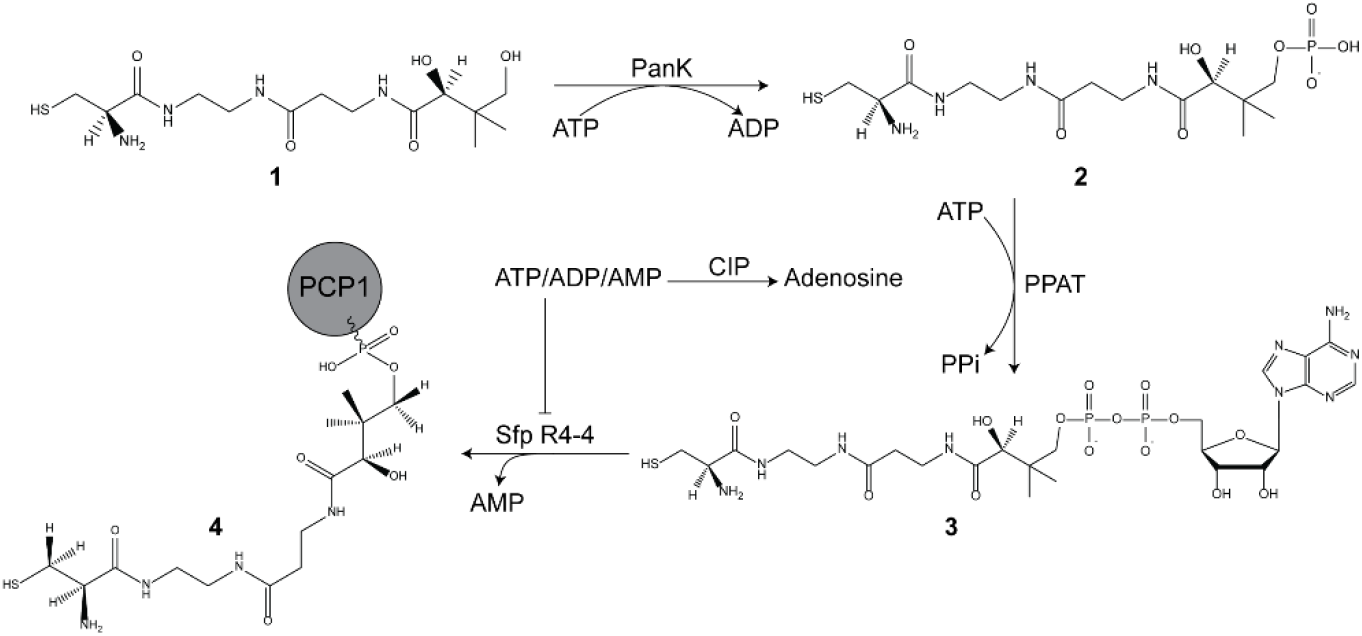
Chemoenzymatic loading of pantetheine analogs to PCP1. A modified one-pot reaction adapted from [11] and [12] is used to generate cysteine-loaded PCP1. A non-hydrolyzable, cysteine-linked pantetheine precursor **1** is modified over a series of sequential reaction steps mediated by the enzymes PanK, PPAT, and Sfp R4-4, with CIP preventing stalling of Sfp R4-4.

Here, we present an NMR framework to monitor each step of the chemoenzymatic reaction and load the peptidyl carrier protein 1 (PCP1) of yersiniabactin synthetase[23] with a stable cysteine-linked pantetheine derivative. We first exploit a differential labeling scheme wherein ^15^N/^13^C labeled apo PCP1 is modified with an unlabeled phosphopantetheine arm harboring a cysteine substrate. We show that each step of the one-pot reaction can be monitored through appropriate pulse sequences, and we present a novel pulse sequence to monitor the final loading step of the reaction. This pulse sequence employs an improved combination of diffusion and isotope filters to isolate the signals of an unlabeled moiety tethered to a labeled protein in presence of a complex mixture of unlabeled reagents, substrates, and small molecule intermediates. We also offer a solution to monitor reactions for routine productions of modified unlabeled proteins. Our approach provides an easy and reliable means to identify experimental errors, defective enzymes, or substrate depletion, such that rescue plans may be designed, and it provides a means to verify that interventions lead to successful protein modification. We illustrate the method both with an uneventful reaction and in a situation in which multiple interventions were necessary.

## Results and Discussion

We first established a protocol to monitor the modification of costly doubly labeled ^15^N/^13^C carrier proteins. Our objective is to employ 1D ^1^H-NMR to rapidly assess the progress of the reaction and not impose delays in incubation times due to data acquisition. Here, we employ a stepwise one-pot protocol, in which the NMR tube initially contains unlabeled pantetheine precursor **1**, small molecule buffer components, ATP, as well as doubly-labeled PCP1 (see Materials and Methods). The pantetheine precursor is typically added in excess of the protein, with production costs limiting the amount, and we often use a 1.5-fold excess with a protein concentration of 100 µM (a value often used in protocols, with 50 µM being otherwise most often used). Here, we used a 5-fold excess to test the method presented in the next paragraph under an extreme case. Each step in the one-pot reaction uniquely modifies the terminal end of the pantetheine substrate, which is converted from a hydroxyl to a phosphoryl group, and to an activated AMP intermediate. As a result, chemical moieties near this modified site may have unique spectral signatures for each intermediate, allowing us to track each step in the reaction up to the addition of CIP. To isolate the signals of unlabeled reagents and intermediates from those of the protein, we used our recently developed 1D isotope filter,[24] consisting of two sequential, tuned isotope X half filters[25–27] targeting methyl (J_CH_=120 Hz) and aliphatic (J_CH_=150 Hz) signals of PCP1, with an additional filter using delayed decoupling to target surviving aromatic signals (J_CH_=220 Hz). We observe that successive modifications of pantetheine derivatives can be monitored through intense signals of methyl groups, providing a convenient readout to estimate the completeness of each step (Figure 2). In this reaction run, PanK converted **1** to **2** with ∼ 100% completion (Figure 2b-c), while the conversion of **2** to **3** (Figure 2c-d) only proceeded to 83% after 1-hour of incubation with PPAT (see also SI Figure 1 for a discussion of PPAT). As each enzyme in the reaction has varying turnover rates, some steps may require longer incubation times, an aspect we will further discuss, and our approach enables to monitor the conversion of each substrate and intermediate to verify that the next step can be performed. CIP is next added to remove phosphoadenylates, and we monitor its activity by inspecting the signals of H8 and H2 groups in adenosine derivatives (see Figure 5 and SI Figure 1). This time a simple 1D NOESY spectrum provides an adequate readout as the concentration of ATP is 2 mM. In the end, we could monitor the multi-step conversion of a derivative of phosphopantetheine to the corresponding dpCoA derivative, but we required a novel pulse sequence to monitor the final attachment of the dpCoA intermediate **3** to PCP1.

**Figure 2.**
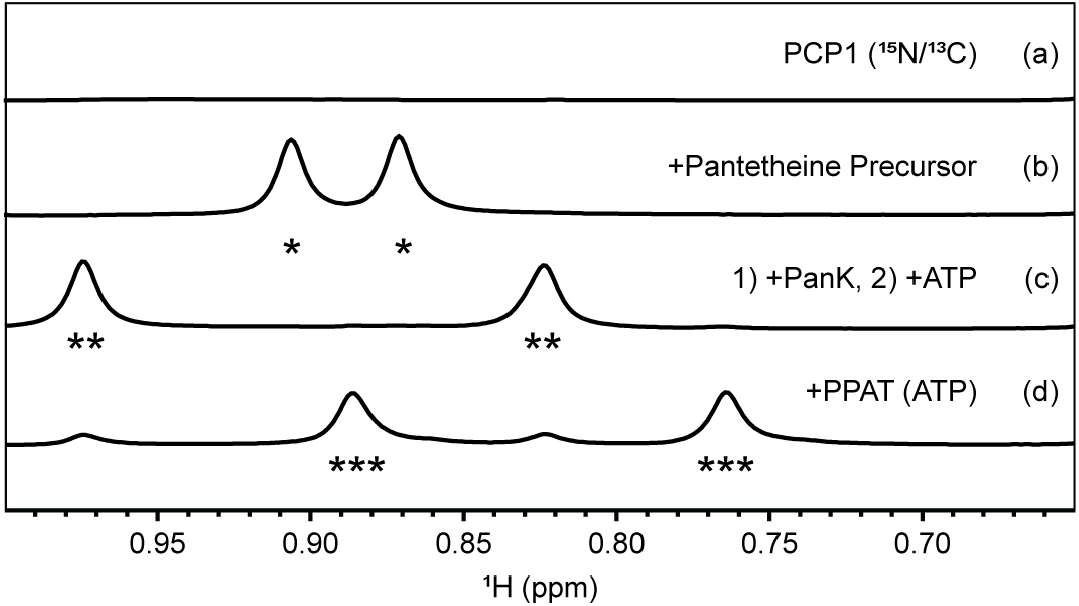
Monitoring pantetheine modifications and reaction completeness through 1D isotope filtered NMR. Each step of the chemoenzymatic modification of an unlabeled cysteine-linked pantetheine precursor was monitored using 1D isotope filtered NMR. (a) Signals of PCP1 were filtered using a sequential tuned X half-filtered pulse sequence (tuned to target 120 and 150 Hz) with delayed decoupling during acquisition for further suppression of aromatic signals (220 Hz).[24] Modification of the starting substrate is conveniently assessed through the intense methyl signals of the moiety. (b) Asterisks (*) denote the two methyl signals of **1**, (c) double asterisks (**) denote the methyl groups of **2**, and (d) triple asterisks (***) denote methyl groups of **3**.

We designed an improved pulse sequence to isolate the signals of an unlabeled moiety tethered to a labeled protein within a solution of complex mixtures of unlabeled reagents. We needed a 1D NMR method that allowed us to monitor the final step in the one-pot reaction, where Sfp R4-4 catalyzes the attachment of the phosphopantetheine group of **3** harboring a cysteine substrate to generate **4**, ^15^N/^13^C loaded PCP1 (Figure 1). Here, an important objective is to demonstrate that new signals appearing in NMR spectra denote loading and not an undesired side reaction leading to untethered pantetheine derivatives. To this aim, we exploited a diffusion filter, the primary building block of Diffusion NMR spectroscopy (DOSY),[28–30] to attenuate signals of fast diffusing reagents and buffer molecules, and we incorporated isotope filters to eliminate signals of the protein core, thereby highlighting the spectrum of the unlabeled moiety covalently attached to the protein. The pulse sequence can be seen as an update on the low-pass isotope filtered diffusion experiment of Gonnella and Co.,[31] now incorporating sequential, tuned isotope filters and our recently developed delayed decoupling filter.[24] Briefly, magnetization is spatially encoded (Figure 3, point A) and decoded (Figure 3, point B) around a diffusion period Δ, such that signals are discriminated against the diffusion of their respective molecules, as described by the Stejskal-Tanner equation[28]:

**Figure 3.**
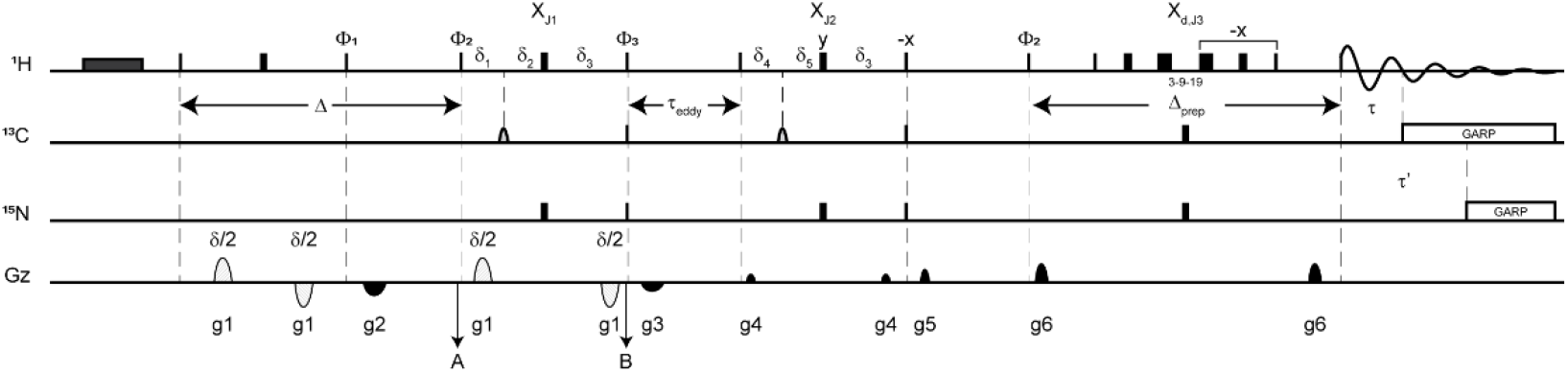
1D diffusion edited, isotope filtered pulse sequence to monitor unlabeled moieties attached to labeled biomolecules. We modified a DOSY sequence using bipolar gradients (dashed half-ellipses) flanking a diffusion delay to incorporate isotope filters. A first isotope X half-filter (X_J1_) is concatenated within the decoding period followed by a sequential tuned X half-filter block (X_J2_) and a shared isotope filtered/Watergate element (X_d,J3_) with delayed decoupling during acquisition.[24] The original Bruker pulse sequence ledbpgp2s1d [32] was first modified as in [33]. The presaturation period (dark grey rectangle) is optional. We used a longitudinal eddy current delay (τ_eddy_) of 5 ms. Narrow and thick rectangles correspond to high power 90° and 180° pulses, respectively. Light gray half ellipses on the carbon channel correspond to 180° frequency-swept chirp inversion pulses, with durations of 500 μs using Bruker shaped pulse Crp60, 0.5, 20.1.[34] Diffusion editing is performed over the period Δ. To detect the phosphopantetheine moiety attached to PCP1, we used a diffusion period (Δ) of 150 ms, encoding/decoding gradient lengths of total duration δ = 2 ms with 47.5 G cm^-1^ strengths. For comparison, reference spectra were obtained with gradient strengths of 2.5 G cm^-1^. The delays during filter blocks X_J1_ and X_J2_ were fixed to target J_NH_ = 90 Hz and J_CH_ values of 120 (methyl) and 150 Hz (aliphatic) signals, respectively, and a filter with delayed decoupling targeting aromatic signals. To accommodate a larger range of decoding gradient lengths, the concatenated filter period X_J1_ should target the smallest J_CH_ coupling constant. The delays in the filter periods are δ_3_ = 1/|4J_NH_|, δ_1_ = 1/|4J_CH,1_|, δ_2_ = 1/|4J_NH_| - 1/|4J_CH,1_|, δ_4_ = 1/|4J_CH,2_|, δ_5_ = 1/|4J_NH_| - 1/|4J_CH, 2_|, with the delayed decoupling filter element (X_d,J3_) designed to target Δ_prep_ + τ’ = 1/|4J_NH_| and Δ_prep_ + τ = 1/|4J_CH,3_|. All filter elements are conveniently coded through constants to target individual coupling constants. Decoupling on carbon and nitrogen is applied with a GARP sequence [35] applied on resonance with the targeted signals using field strengths of 2.0833 kHz and 1.042 kHz, respectively. The phase cycle used was ϕ_1_ = x, x, -x, -x, ϕ_2_ = x, x, x, x, -x, -x, -x, -x, ϕ_3_ = x, -x, x, -x, -x, x, -x, x, and ϕ_rec_ = x, -x, -x, x, -x, x, x, -x, with all remaining phases being applied along x. Gradients were applied with lengths (τg_i_) and strengths (g_i_) along the z-axis with τg_1_ = δ/2 = 1 ms (bipolar gradients, Bruker shape SMSQ10.32), τg_5_ = τg_6_ = 1 ms (Bruker shape SMSQ10.32), τg_2_ = τg_3_ = = τg_4_ = 300 µs (Bruker shape SMSQ10.32), g_1_ = 47.5 G cm^-1^, g_2_ = 8.565 G cm^-1^, g_3_ = 6.585 G cm^-1^, g_4_ = 2.5 G cm^-1^, g_5_ =5.5 G cm^-1^, and g_6_ = 10 G cm^-1^.

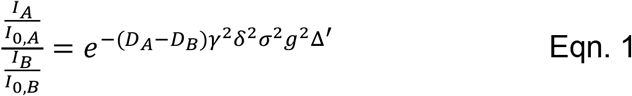

where I_i_ is the intensity of a signal for molecule I diffusing during Δ, I_0,i_ is the intensity of a signal unattenuated by diffusion, D_i_ is the diffusion coefficient, γ is the gyromagnetic ratio of the nuclei, δ is the length of the encoding and decoding gradients, σ is the gradient shape factor, g is the strength of the encoding/decoding gradients, and Δ’ is the duration of the diffusion period corrected to account for the shape of pulsed-field gradients. Thus, signals of slowly diffusing molecules are less attenuated at the end of Δ’ than those of fast diffusing molecules, e.g. reagents and buffer components, with the difference between diffusion coefficients defining the resulting discrimination. We next sought to eliminate signals of the protein core through isotope filters. To mitigate losses due to relaxation, the first isotope half-filter X_J1_ is concatenated with the diffusion decoding period. Thus, following the decoding period, the coherences of protons attached to isotopically active heteroatoms are eliminated by purge gradients, while those of the unlabeled moiety of interest are preserved. Concatenation is not strictly necessary for proteins as small as PCP1 (∼10 kDa), where only 2.2% of losses are expected, but was implemented for broader application of the pulse sequence to larger multidomain constructs. A second tuned isotope half filter and an additional filter using delayed decoupling further suppress signals of the labeled protein as described in [24]. We set our three isotope filters to target methyl, X_J1_, aliphatic, X_J2_, and aromatic, X_J3,d_ moieties. It is important to target the moiety featuring the smallest scalar coupling, here 125 Hz, during the filter concatenated with the diffusion decoding period, X_J1_, so that long decoding gradients may be employed. In the end, this 1D isotope diffusion filtered experiment unambiguously identifies signals of the attached unlabeled moieties (Figure 4). With this suite of experiments, we were able to monitor the entire chemoenzymatic conversion of a modified pantetheine analog including its attachment to doubly labeled PCP1. We note that the objective is to verify that the reaction proceeds as expected and not to provide an overall yield of conversion. The final sample conversion yield is assessed through MALDI-TOF-MS or, if possible, a 2D HN-HSQC (SI Figure 4), or both.

**Figure 4.**
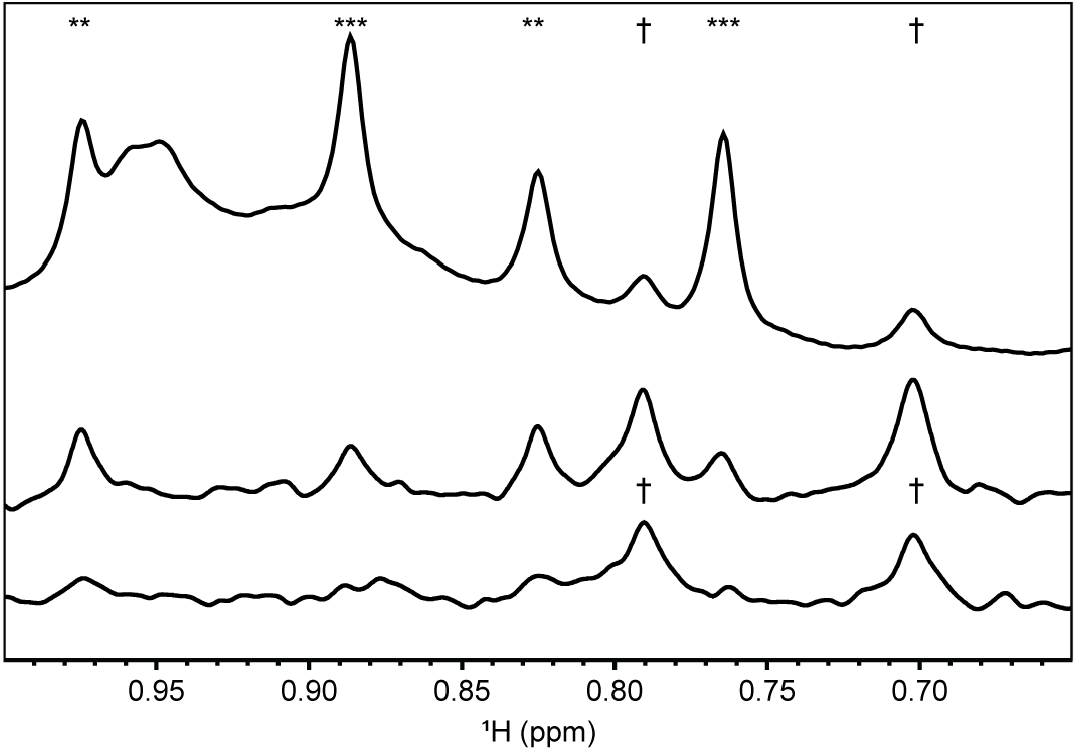
An isotope and diffusion filtered 1D experiment identifies signals of unlabeled moieties attached to labeled proteins. We utilized a novel, sequential, tuned isotope and diffusion filtered pulse sequence (Figure 3) with delayed decoupling to selectively observe methyl groups from the unlabeled phosphopantetheine moiety of **3** (Figure 1) attached to _15_N/_13_C labeled PCP1 in presence of unlabeled reagents (bottom). The diffusion period was set to 150 ms and the bipolar gradient strength to 47.5 G cm_-1_. The same experiment with gradient strengths at 2.5 G cm_-1_ is shown in the middle to highlight the suppression of signals of fast-diffusing free substrates. A conventional 1D is also shown at the top. Double asterisks (**) denote the signals of methyls in **2**, and triple asterisks (***) denote those in **3**. † denote the signals of methyl groups upon covalent attachment of the phosphopantetheine arm to PCP1 (**4**). The preservation of these signals through a diffusion filter demonstrates that they belong to a moiety attached to the protein.

This pair of isotope edited experiments not only provided a means to follow the reaction when modifying a labeled protein, with applications illustrated below, but it also permitted to test a readout for monitoring the conversion of unlabeled carrier proteins. Given that we have unambiguously identified NMR readouts of all steps in the reaction with the labeled protein (Figures 2 and 4), we next investigated what experiments provided the best readouts when modifying an unlabeled protein. Here, diffusion and T2 filters can be exploited to emphasize signals of interest. Thus, the first steps of the one-pot reaction, preceding loading, can all be monitored through 1D T2 filters to attenuate the signals of the protein core (SI Figure 2a,b). We cannot isolate signals of the loaded moiety in the last step but can nevertheless identify them by comparing diffusion edited spectra (SI Figure 2 c,d). Although the last step is only assessed in a very qualitative manner, verifying that the preceding steps have gone to completion and that loading is detected is nevertheless a powerful advantage as rescue plans may be designed on the spot as illustrated below for a doubly labeled protein.

Monitoring chemoenzymatic reactions by NMR permits to identify pitfalls and verify that remediations are successful. While we presented our methodology using the step-wise protocol described by Cryle and Co., we have also monitored reactions when we collected data to design, implement, and optimize novel pulse sequences. Under these conditions, timing could not be followed as the protein was incubated with enzymes for hours (see Methods) rather than 5 to 30 minutes as implemented by Cryle and Co. Most notably, when first optimizing the diffusion isotope edited pulse sequence, all previous steps had been subject to incubations longer than usual, and the sample was left to incubate overnight after addition of Sfp R4-4 at a late hour. We not only found that loading had stalled to 41% (Figure 5a) but also that signals of molecules in the preceding steps were visible in the spectrum, including the original starting material, **1**. Looking at data at the beginning and end of the incubation revealed that phosphoderivatives of adenosine were still present before incubation but all converted to adenosine 10 hours later (Figure 5b,c), so stalling was not due to poisoning of Sfp R4-4. We concluded that CIP had outcompeted Sfp R4-4, most likely because the concentration of Sfp R4-4 was lower than expected from calculations, and that the remaining enzymes were then favoring the reverse reactions in absence of ATP, leading to **1**. We next tried to rescue the reaction by adding more ATP, and we could verify the conversion of **1** to **2** and to the dpCoA derivative **3**, before adding CIP and Sfp R4-4, the latter now being in excess. Conversion of phosphoderivatives was again monitored (Figure 5 e, f), and this time we obtained conversion to 87 % (Figure 5d), and 100% of conversion was reached after spiking with ATP and precursors. The details of concentrations and timing are reported in the Method section. Clearly, these rather desperate measures were the consequences of highly inappropriate timing when first converting the pantetheine derivative to its dpCoA counterpart in the first place. Nevertheless, similar situations may occur accidentally during routine reactions. For example, reagents may be contaminated or the concentration of enzymes in the reaction tubes may differ from that predicted from the stock solutions, e.g. because of degradation or precipitation in the thawing stocks or because of mis-calibrated pipettors. Although SDS-PAGE or MALDI-TOF-MS would be needed to confirm these hypotheses and provide a long-term remediation, such as changing reagents or measuring the enzyme concentration in stock solutions, monitoring the reaction by NMR immediately identifies issues and allows for rescuing on the spot. For example, in a different reaction, we identified that ATP had been depleted when adding the second enzyme, PPAT, and addition of ATP rescued the reaction that had stalled at that stage (data not shown). We conclude this section by highlighting that, although we have employed the sequential one-pot protocol to be able to set-up novel experiments, the all in-one approach that we also routinely employ can be monitored through the same experiments. Notably, the spectrum of CoA is markedly different from that of dpCoA (SI Figure 3), and users who favor using the enzyme DPCK and wild-type Sfp may use our approach. In closing, we believe that our NMR framework is a formidable tool to prevent costly and time-consuming repeats in sample preparation and to provide homogeneous samples with fully modified proteins, thereby enhancing the reliability and reproducibility of subsequent experiments.

**Figure 5.**
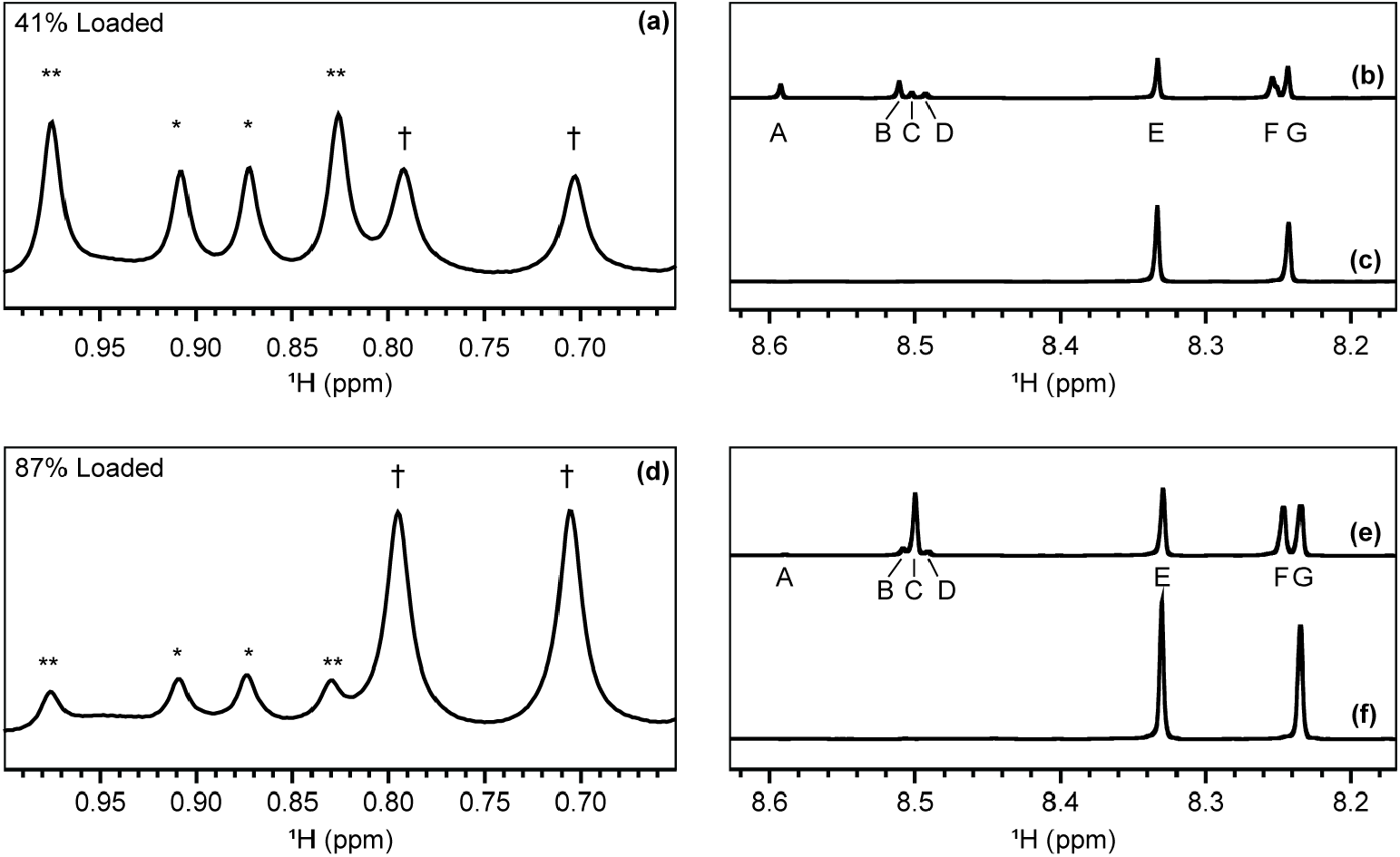
Debugging and rescuing the complex chemoenzymatic reaction with 1D NMR. The 1D isotope filtered spectrum (a) showed that loading only proceeded to 41% (signals †) and that the precursor **3** was converted back to **2** (**) and the starting material **1** (*). The downfield regions of 1D NOESY spectra before (b) and after overnight incubation (c) demonstrate that CIP converted all phosphoadenylates, AMP (signal A), ADP (B), and ATP (C) to adenosine (signals E and G). To rescue the reaction ATP, CIP, and Sfp R4-4 were supplemented to the reaction, and the increase in Sfp R4-4 concentration led to 87% of loaded PCP1 conversion (d). Spectra before (e) and after final incubation (f) indicated that all the ATP in the reaction had been dephosphorylated by CIP, as needed to prevent Sfp R4-4 inhibition. A final addition of ATP and starting material **1** led to 100% conversion under the final enzyme concentrations. See Material and Methods for details.

## Conclusion

We have provided a 1D NMR framework to monitor complex chemoenzymatic reactions that modify proteins with an unlabeled moiety. The method was designed to follow a one-pot reaction involving four enzymes, which successively modify derivatives of pantetheines and tether them to a carrier protein. For a labelled protein, we employ 1D isotope filters to monitor modifications of the unattached precursors. We presented a novel diffusion isotope filtered experiment to isolate signals of the unlabeled moiety tethered to the labeled protein within a background of unlabeled reagents. For an unlabeled protein, the steps preceding loading can all be followed with T2 filters, and loading can be observed through a diffusion filtered 1D experiment. These 1D NMR readouts benefit from unambiguous spectral signatures of chemical modifications affecting intense methyl signals. This methodology enabled us not only to follow the progress of a successful reaction but also to rescue a reaction that had been stalled. The method readily identifies when enzymes are less active, substrates are limiting, or more generally when reaction kinetics impede product formation, e.g. when an undesirable reverse reaction dominates. This approach will facilitate otherwise cumbersome reaction optimizations that provide homogenous samples and enhance the reproducibility and integrity of subsequent studies by NMR or other techniques. Our study illustrates the critical role that NMR may play in enhancing the scientific rigor and reproducibility of biomolecular studies due to its non-invasive nature and an exquisite sensitivity to molecular changes.

## Materials and Methods

### Isotope labeling, expression, and purification of PCP1

Doubly labeled ^15^N/^13^C apo PCP1 of the *Yersinia pestis* irp2 1402-1482 gene fragment (accession number AAM85957) was expressed and purified as described in [24] in an *E. coli* BL21 (DE3) ΔEntD cell line (courtesy of Christian Chalut and Christophe Guilhot, CNRS, Toulouse, France). Isotope labeling was performed using M9 minimal media containing 1 g L^-1 15^NH_4_Cl and 2 g L^-1 13^C glucose to generate uniformly labeled ^15^N/^13^C PCP1.

Expression and purification of one-pot enzymes PanK, PPAT, and Sfp R4-4 were produced as described in [16] (PanK, PPAT) and [36] (Sfp R4-4).

Isotopes used in the labeling of PCP1, buffer reagents used in the purification, and Quick CIP (alkaline phosphatase) were purchased from Cambridge Isotope Laboratories, VWR, and New England Biolabs, respectively.

### Synthesis of an unlabeled non-hydrolyzable cysteine-linked pantetheine precursor

Loading of apo ^15^N/^13^C apo PCP1 was performed using a cysteine-linked pantetheine precursor wherein the native thioester linkage was replaced with an amino group to overcome hydrolysis of the cysteine substrate in solution (SI Figure 5). The synthesis is described below.

Synthesis of **III**: Boc-Cys(Trt)-OH **I** (1466 mg, 3.16 mmol) was mixed with HATU (1732 mg, 4.55 mmol) and DIEA (793 ul, 4.55 mmol) in DMF (12 ml) at room temperature for 10 minutes. The activated mixture was added to amine **II** (prepared according to [37]) in DMF (12 ml) and the resulting mixture was stirred for 12 hours. The reaction was quenched with water (60 ml) and the aqueous phase was extracted with EtOAc (60 ml x3). The combined organic phase was washed with 0.5 N aq. HCl (60 ml), water (60 ml), saturated NaHCO_3_ (60 ml), brine (60 ml x3), dried with anhydrous Na_2_SO_4_, and concentrated under reduced pressure. The concentrated crude product was purified by flash column chromatography (0 to 10% MeOH in CH_2_Cl_2_) to give **III** (1052 mg, 48%). ^1^H NMR (500 MHz, CDCl_3_) δ 7.42-7.47 (m, 8H), 7.30-7.34 (m, 8H), 7.26 (d, *J*= 8.7 Hz, 2H), 7.01 (br, 1H), 6.94 (d, *J*= 8.7 Hz), 2H), 6.58 (br, 1H), 6.50 (br, 1H), 5.50 (s, 1H), 4.94 (br, 1H), 4.12 (s, 1H), 3.84 (s, 3H), 3.83-3.88 (m, 1H), 3.71 (dd, *J1*= 11.3 Hz, *J2*= 25.6 Hz, 2H), 3.23-3.58 (m, 6H), 2.75 (dd, *J1*= 6.8 Hz, *J2*= 12.7 Hz, 1H), 2.55 (dd, *J1*= 5.7 Hz, *J2*= 12.7 Hz, 1H), 2.30 (br, t, *J*= 7.5 Hz, 2H) 1.44 (s, 9H), 1,15 (s, 3H), 1.12 (s, 3H). ^13^C NMR (126 MHz, CDCl_3_) δ 171.33, 171.23, 169.45, 160.20, 144.23, 130.14, 129.49, 128.07, 126.95, 113.70, 101.33, 83.84, 78.45, 67.20, 55.30, 39.29, 36.21, 34.94, 33.06, 28.27, 21.86, 19.12.

Synthesis of **IV**: To a solution of compound **III** in THF (20 ml), 1 N aqueous HCl (20 ml) was added slowly at room temperature and stirred for one hour. The resulting solution was neutralized by adding 1 N NaOH (10 ml) and saturated NaHCO_3_ (7.5 ml) and extracted with EtOAc. The combined extract was washed with water, brine (2x), and dried with Na_2_SO_4_. Purification by flash column chromatography (0 to 15% MeOH in methylene chloride) gave diol **IV** (592 mg, 69%) as a white solid. ^1^H NMR (500 MHz, CDCl_3_) δ 7.39-7.47 (m, 6H), 7.29-7.36 (m, 6H), 7.24-7.28 (m, 2H), 6.96 (br, 1H), 6.85 (br, 1H), 5.18 (br, 1H), 4,81 (d, *J*= 5.2 Hz, 1H), 4.09 (br, s, 1H), 4.01 (d, *J*= 5.0 Hz, 1H), 3.90 (dd, *J1*= 6.4 Hz, *J2*= 12.9 Hz, 1H),3.58 (br, s, 1H), 3.41-3.52 (m, 3H), 3.21-3.41 (m, 4H), 2.72 (dd, *J1*= 6.9 Hz, *J2*= 12.7 Hz, 1H), 2.56 (dd, *J1*= 5.3 Hz, *J2*= 12.7 Hz, 1H) 2.56 (m, 1H), 2.32 (br, 1H), 1.44 (s, 9H), 1.02 (s, 3H), 0.94 (s, 3H).

Synthesis of **Cys-NH-Pantetheine analog**: To the diol **IV** (100 mg, 0.14 mmol) was added a cleavage cocktail (5 ml, TIPS/TFA = 5/95 (v/v)). Incubation was done at room temperature for 5 min and stopped by the addition of cold diethyl ether (40 ml). The precipitated white solid was centrifuged down, and the supernatant was decanted. The ether wash/centrifugation/decant steps were repeated 3 times in total. The final white precipitate was dried under N_2_ and collected as a white solid (**Cys-NH-Pantetheine**, 40 mg, 78%), which was used for the next enzymatic assays without any further purification steps. ^1^H NMR (500 MHz, CD_3_OD) δ 3.99 (dd, *J1*= 5.0 Hz, *J2*= 6.9 Hz, 1H), 3.92 (s, 1H), 3.45-3.55 (m, 3H), 3.28-3.44 (m, 5H), 3.07 (dd, *J1*= 5.0 Hz, *J2*= 14.6 Hz, 1H), 2.96 (dd, *J1*= 7.1 Hz, *J2*= 14.6 Hz, 1H), 2.43 (t, *J*= 6.6 Hz, 1H), 0.94 (s, 6H). ^13^C NMR (126 MHz, CD_3_OD) δ 215.67, 174.56, 168.74, 77.46, 70.42, 56.41, 40.72, 39.92, 37.08, 36.61, 26.34, 21.54, 21.03.

### Chemoenzymatic loading of doubly labeled PCP1 with an unlabeled pantetheine analog

We implemented the one-pot reaction (Figure 1) in a stepwise manner to monitor each step towards loading of PCP1. All reaction steps were performed at 25 °C and NMR experiments were optimized and collected at each step upon addition of enzymes and/or reagents. ATP was prepared as a 100 mM stock in one-pot reaction buffer (100 mM Tris, pH 7.55 at 22 °C, 10 mM MgCl_2_, 100 mM NaCl, 2.5 mM DTT, 10% v/v D_2_O, and 200 µM DSS for referencing). We used 200 µM stock solutions of PanK and PPAT in 20 mM Tris pH 7.5, 100 mM NaCl, and 10% (v/v) glycerol. We used a 2.5 mM stock solution of Sfp R4-4 in 10 mM Tris pH 7.5, 1 mM EDTA, and supplemented with 10% glycerol. Quick CIP stocks were used at 500 U mL^-1^ in one-pot reaction buffer.

In a first reaction providing the data for Figures 2 and 4, 500 μL of ^15^N/^13^C labeled apo PCP1 (100.1 µM) in one-pot reaction buffer was transferred to an NMR tube and degassed. Reference 1D NOESY, 1D isotope filter, 1D isotope and diffusion filtered, and 2D HN-HSQC experiments were collected. The sample was supplemented with 473.5 µM of cysteine-linked pantetheine precursor **1** (from a stock at 7.62 mM in water). PanK was added to 2.5 µM, and reference spectra were again collected for 42 minutes. The reaction was initiated with the addition of ATP (2.3 mM) with an incubation of 35 minutes during which more spectra were recorded. PPAT was then added to 2.5 µM, and the reaction was incubated for approximately one hour as NMR spectra were collected. Quick CIP (1.9 U mL^-1^) was supplemented into the reaction and incubated for one hour and 38 minutes. Sfp R4-4 was added to 2.4 µM and incubated for two hours and 33 minutes. Following this period, apo PCP1 was fully converted to loaded PCP1 as determined by 2D HN-HSQC spectra, where loading efficacy was determined to be 100%. Accounting for all dilutions, the final sample contained 94.3 µM PCP1, 2.4 µM PanK, 2.4 µM PPAT, 1.9 U mL^-1^ Quick CIP, 2.4 µM Sfp R4-4, 2.4 mM ATP, and 471.2 µM Cys-NH-pantetheine analog. We underline that the acquisition of 1D isotope filtered, 1D-NOESY, and 1D isotope and diffusion filtered spectra each only take 2, 3, and 2 minutes, respectively.

We repeated this stepwise, one-pot reaction on a larger scale (∼5 mL) but faced issues with incomplete loading of PCP1 due to prolonged incubation steps. This reaction was ultimately rescued and provided the data for Figure 5 in the main text. At the start, ^15^N/^13^C apo PCP1 (134.3 µM) was supplemented with 544.1 µM of the cysteine-linked pantetheine precursor **1** in one-pot reaction buffer containing 10% (v/v) D_2_O and DSS (200 µM). PanK (2.6 µM) was added and incubated at 25 °C for 50 minutes. The reaction was started with the addition of ATP (2.6 mM) and incubated for 51 minutes. PPAT (2.5 µM) was then added and the reaction was incubated for one hour and two minutes. To assess whether Sfp R4-4 could efficiently load with phosphoadenylates present, we next added 2.5 µM Sfp R4-4 and incubated for one hour and 33 minutes. Only 5% loading of PCP1 was observed consistent with observations made by Cryle and Co.[12] Next, Quick CIP (2.0 U mL^-1^) was added and incubated for 1 hour and 40 minutes. Additional Sfp R4-4 (2.5 µM) was then added to try to push the reaction to completion. Following overnight incubation for 9 hours and 51 minutes, PCP1 still showed incomplete loading. To restart the reaction as the original pantetheine analog **1** had been reformed, ATP (2.5 mM) was added to the reaction and incubated for one hour and 43 minutes. Next, Quick CIP (0.5 U mL^-1^) was supplemented into the reaction and incubated for 43 minutes. An additional 4.9 µM Sfp R4-4 was then added to favor loading in presence of CIP. The reaction was incubated for 10 hours and 12 minutes overnight. Following overnight incubation, the reaction was supplemented with **1** (300.6 uM) and ATP (2.3 mM). The reaction was incubated for 1.5 hours after which PCP1 was determined to be fully loaded upon inspection of a 2D HN-HSQC spectrum. Accounting for all dilutions, the final reaction contained 106.4 µM PCP1, 2.3 µM PanK, 2.3 µM PPAT, 2.3 U mL^-1^ Quick CIP, 9.3 µM Sfp R4-4, 7.0 mM adenosine derivatives (ATP/ADP/AMP, and adenosine as a product of CIP), and 757.5 µM Cys-NH-pantetheine analog. The prolonged incubation times resulted from attempts at setting up new experiments not discussed in the manuscript.

### NMR data acquisition

All NMR experiments were collected at 25 °C on a 600 MHz Bruker Avance III spectrometer equipped with a QCI cryoprobe using samples described in the previous sections. All spectra were processed in TopSpin 3.6.2.

All 1D experiments were collected with 32 scans (except the 1D NOESY which was collected with 16 scans), a spectral width of 16.0192 ppm centered at 4.698 ppm, and a 2 sec recycling delay. All 1D isotope filtered NMR experiments were collected with 3084 complex points. These experiments implement a 3-9-19 WATERGATE element for water suppression. Spectra were zero-filled to 16384 points, referenced to DSS, baseline corrected, and apodized using exponential multiplication (5 Hz) to reduce truncation artifacts from intense buffer signals. 1D NOESY experiments were collected using Bruker pulse program noesygppr1d, with continuous-wave presaturation during the recycling delay (2 seconds) and during the NOE mixing time (150 ms) applied on resonance with water with a field strength of 27.3 Hz. The 1D NOESY spectra were collected with 32,768 complex points, zero-filled to 131072 points, and referenced to DSS. All 1D diffusion and isotope filtered experiments implementing the pulse sequence in Figure 3 were collected with 3084 complex points, zero-filled to 16384 points, apodized with a 4 Hz exponential function, and referenced to DSS. Continuous-wave presaturation was applied during the recycling delay with a field strength of 27.3 Hz. For the 1D diffusion spectra, we modified the Bruker pulse sequence ledbpgp2s1d in accordance with [33] and included ^15^N and ^13^C decoupling during acquisition. 1D diffusion spectra (Bruker pulse sequence ledbpgp2s1d) were collected with the same parameters as their isotope filtered counterparts (Figure 3), but with 128 scans. 1D T2 filters were run using the Bruker pulse sequence cpmgpr1d but modified to include ^15^N and ^13^C decoupling during acquisition. Spectra were collected with a fixed echo time of 2 ms and a T2 filter length of 408 ms to attenuate PCP1 signals, while a 24 ms T2 filter was used for reference spectra. Spectra were collected with 128 scans, zero-filled to 16384 points, apodized with a 5 Hz exponential function, and referenced to DSS.

All 2D HN-HSQC spectra used to quantify loading of PCP1 at the end of the reaction were collected with 8 scans, 1 sec recycling delay, 128 complex points in the ^15^N dimension, 2048 points in the detected dimension, spectral widths of 16.0192 ppm (^1^H) and 28 ppm (^15^N), with carriers set at 4.698 ppm (^1^H) and 117.0 ppm (^15^N). Nitrogen decoupling was applied with 1.042 kHz field strength using a GARP sequence. Spectra were apodized with a cosine-squared bell function, extracted over the amide region (^1^H), zero-filled to 2048 (^1^H) and 512 (^15^N) points, and referenced directly (^1^H) and indirectly (^15^N) using DSS.

The assignment of the cysteine-linked pantetheine analog **1** was aided by existing assignments of pantetheine derivatives[39,40] and completed with the following experiments using a 1 mM sample of **1** in water (pH 5-6), containing 10% (v/v) D_2_O, and 200 µM DSS.A 2D ^1^H-^1^H TOCSY spectrum (Bruker pulse sequence dipsi2esgpph) was collected with 60 ms DIPSI-2 [38] mixing sequence applied with a field strength of 10.006 kHz, 2048 t2 points and 512 t1 complex points, spectral widths of 16.0192 ppm (F2) and 16 ppm (F1), carriers at 4.697 ppm, 1 second recycling delay, and 16 scans. Additionally, a 2D NOESY (Bruker pulse sequence noesyesgpph) spectrum was collected using a 200 ms mixing time, 16 scans, spectral widths of 16.0192 ppm (F1 and F2 dimensions), with all other acquisition parameters as in the 2D TOCSY. To assign carbon resonances of **1**, a 2D HC-HSQC spectrum (Bruker pulse sequence hsqcetgpsisp2) was collected with the following parameters: 16 scans, 1.5 sec recycling delay, 2048 (t2) and 256 (t2) points, spectral widths of 16.0192 ppm (^1^H) and 100 ppm (^13^C), and carriers of 4.698 ppm (^1^H) and 50 ppm (^13^C). Assignment of ^15^N resonances in **1** was performed using a 2D HN-HSQC (Bruker pulse sequence hsqcfpf3gpphwg) using: 256 scans, 1 second recycling delay, 2048 (^1^H) and 128 (^15^N) points, spectral widths of 16.0192 ppm (^1^H) and 28 ppm (^15^N), and carriers of 4.698 ppm (^1^H) and 117 ppm (^15^N). All spectra were referenced with respect to DSS before assigning resonances.

## Author Contributions

D.P.F and K.A.M designed experiments. K.A.M prepared all samples, implemented the pulse sequences, recorded, and analyzed all data. K.A.M and E.S.K assigned resonances. D.J.M and Y.H synthesized pantetheine analog. D.P.F and K.A.M wrote the manuscript with feedback from all co-authors. All authors have read and approved the final manuscript.

## Code Availability

The newly introduced isotope and diffusion filtered 1D NMR pulse sequence is readily available from https://frueh.med.jhmi.edu/software-downloads/.

## Supplementary Information

**SI Figure 1.**
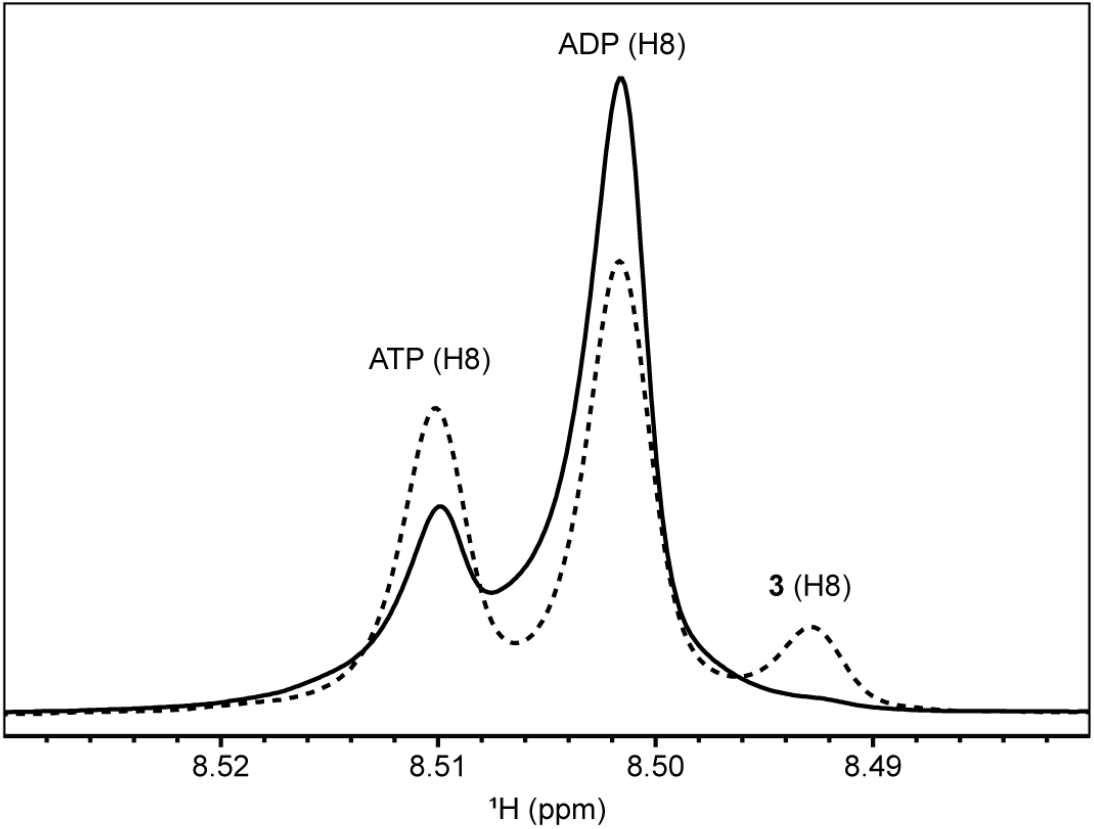
Identifying and overcoming the reversibility of PPAT. PPAT, the enzyme converting **2** to **3** is known to be prone to catalyzing the reverse reaction.[20] To mitigate reductions in yields due to the reverse reaction, we not only monitor the methyl groups of pantetheine derivatives but also signals of phosphoadenylates in 1D NOESY spectra. The spectrum in solid black corresponds to a sample before addition of PPAT, that is where PanK was added and converted **1** into **2** upon addition of ATP and incubation at 25 °C for 50 minutes. At that stage, much of the ATP has been converted to ADP but about ¼ of the starting ATP remains. After addition of PPAT and incubation at 25 °C for one hour (black dashed), we see an increase in ATP, consistent with the reversibility of PPAT. However, signals of **3** are also detected and, in this example, the reaction proceeded to completion. When conversion of **2** to **3** is limited, addition of ATP rescues this step. The signals are those of H8 in ATP, ADP, and **3**.

**SI Figure 2.**
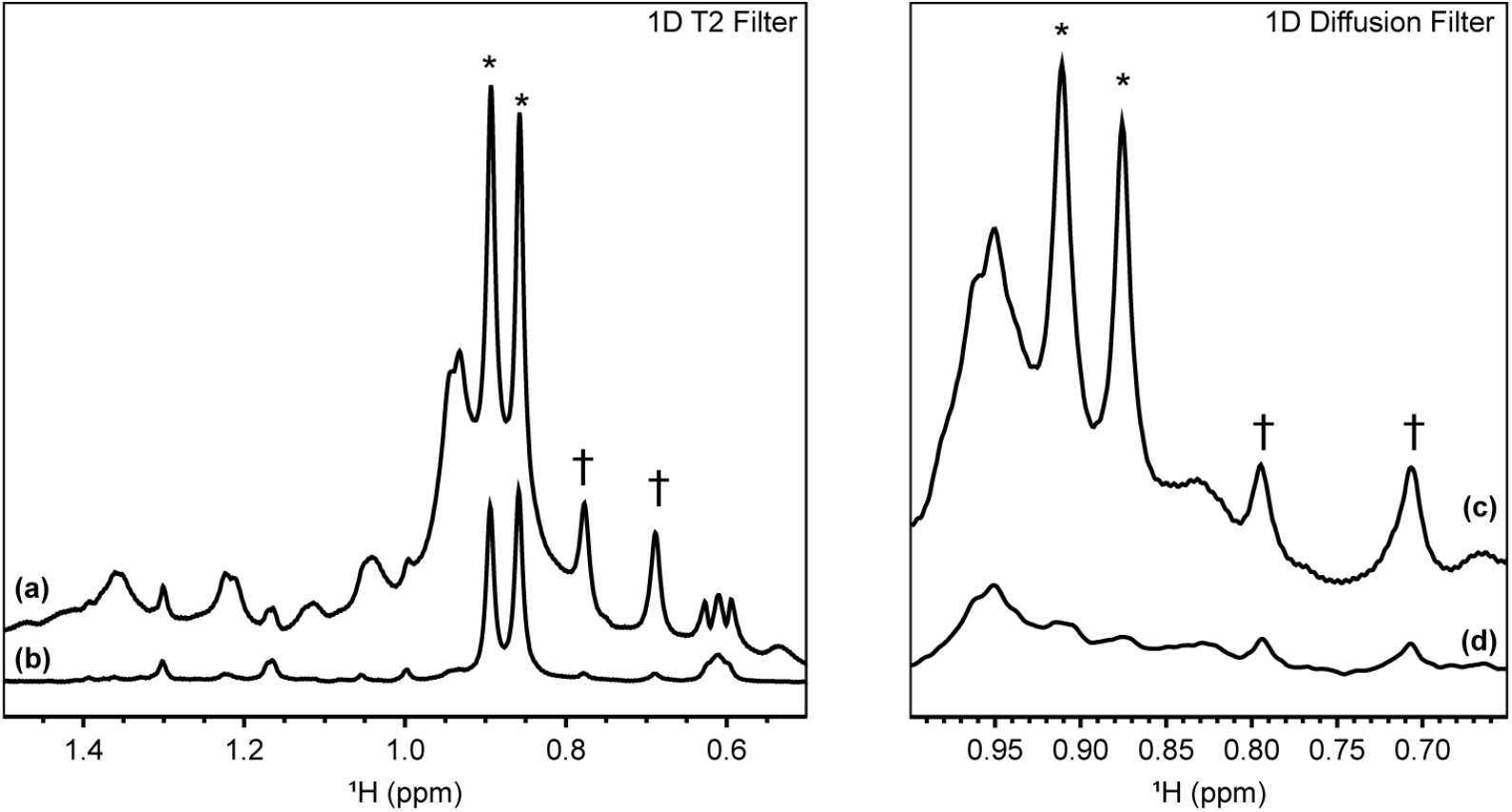
NMR methods for monitoring one-pot reactions of unlabeled proteins. a, b) T2 filters emphasize signals of fast tumbling reagents, albeit with sensitivity losses. T2 filter of 24 ms (a) and 408 ms (b) are shown for comparison.They can be used to monitor all steps preceding loading. c, d) Diffusion filters can be used to identify signals of new methyl groups tethered to a protein as those of precursors are attenuated (*) but not those attached to the protein (†). The spectra were recorded on the doubly labeled sampled used for Figures 2 and 4. ^15^N and ^13^C decoupling was used to emulate an unlabeled sample, and the parameters for the diffusion filter were as described in Figure 3. Asterisks (*) denote the two methyl signals of **1** and † denote the methyl signals upon covalent attachment of the phosphopantetheine arm to PCP1.

**SI Figure 3.**
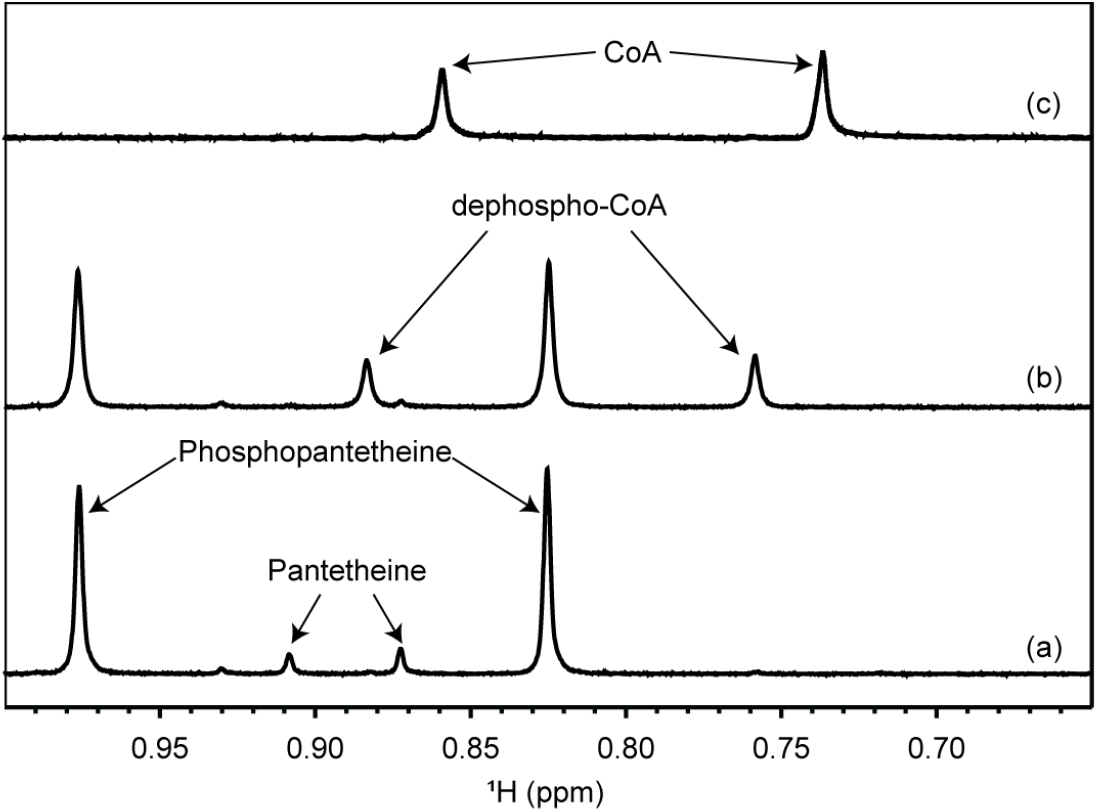
Comparison of methyl signal shifts in pantetheine derivatives. Methyl signals of pantetheine derivates exhibit large perturbations upon modification. 1D NOESY spectra of a sample that converted pantetheine to phosphopantetheine via PanK (a), then to dephospho-CoA via PPAT (b), and a standard sample of CoA (c). All spectra were recorded in one-pot buffer (see Methods) at 25 °C and referenced to DSS. The comparison demonstrates that one-pot reactions relying on DPCK, which generates CoA, can also be monitored with our protocol.

**SI Figure 4.**
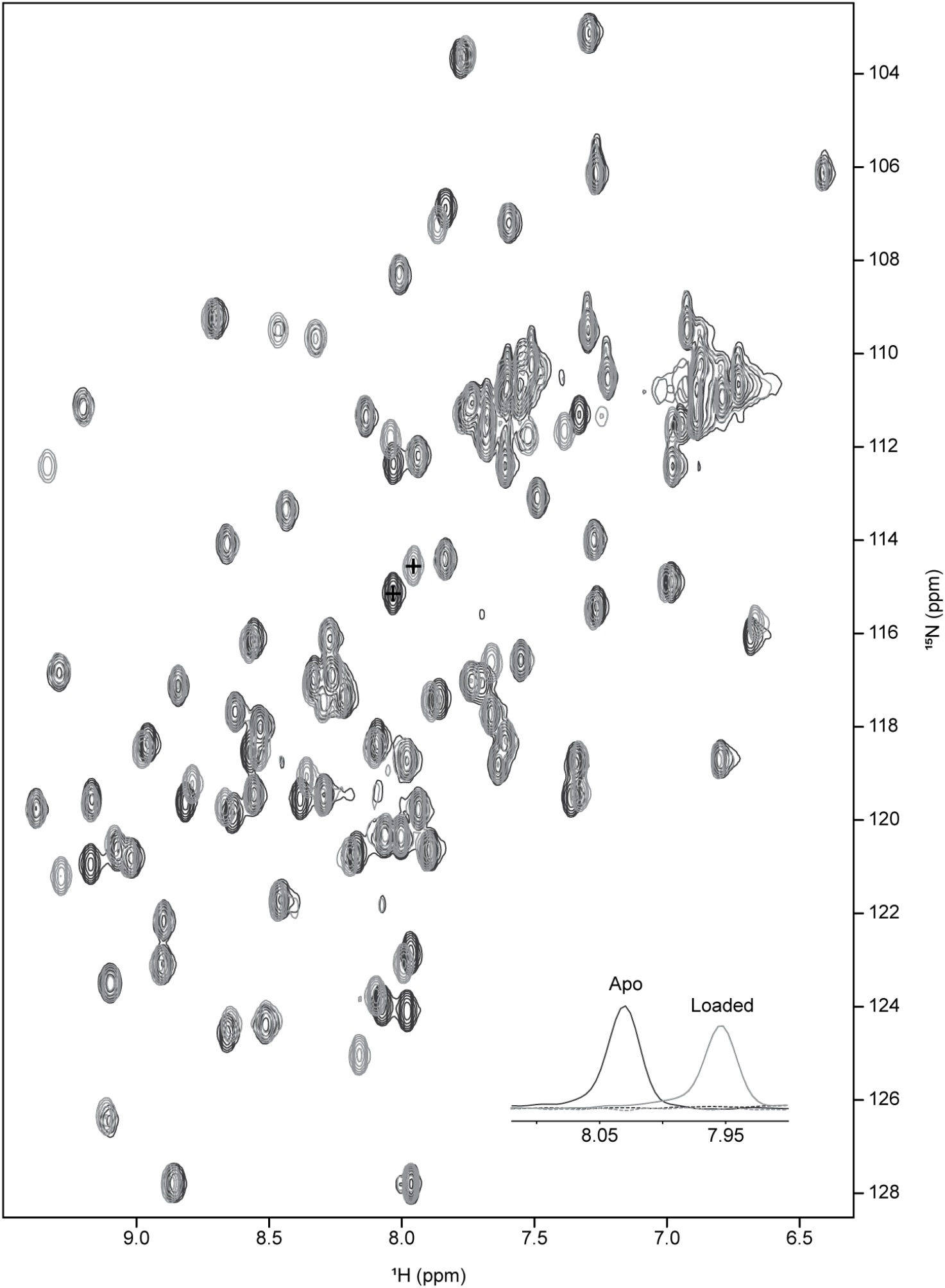
2D HN-HSQC to assess the conversion yield for labeled PCP1. The final conversion yield of the one pot reaction can be assessed through 2D HN-HSQC spectra of apo _15_N/_13_C PCP1 before loading (black) and after loading (light gray). Inset shows 1D slices through signals of apo (black) and loaded PCP1 (light gray) at positions marked with (+). The black dashed spectrum indicates the 1D slice through the apo spectrum at the position of the loaded peak, and the light gray dashed spectrum indicates the 1D slice through the loaded spectrum at the position of the apo peak.

**SI Figure 5.**
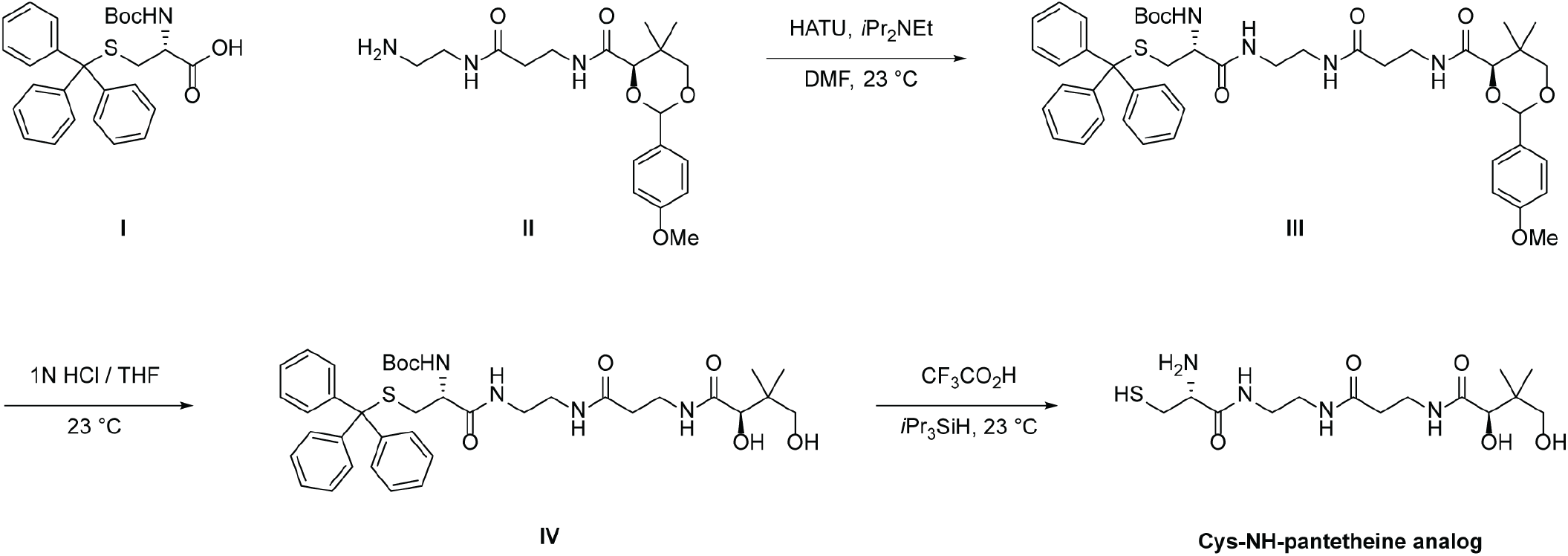
Synthesis of a cysteine-linked, non-hydrolyzable pantetheine analog.

**SI Figure S6.**
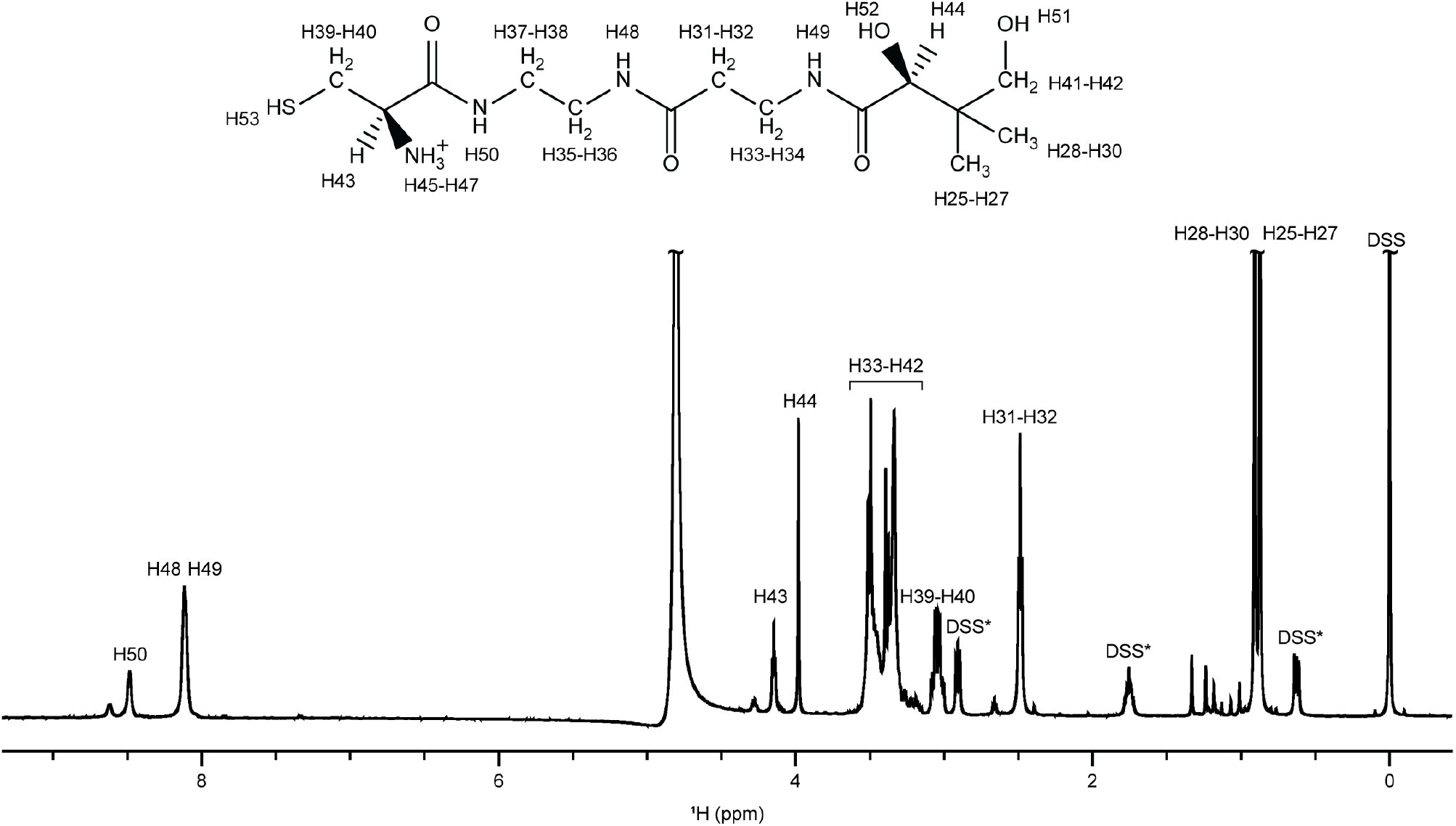
Chemical shift assignment of Cys-NH pantetheine analog 1. 1D spectrum with proton resonances of **1** labeled as defined on the chemical structure. Peaks labelled as DSS* correspond to DSS impurities. Atom labels were generated using ALATIS.[41]

**SI Table 1.**
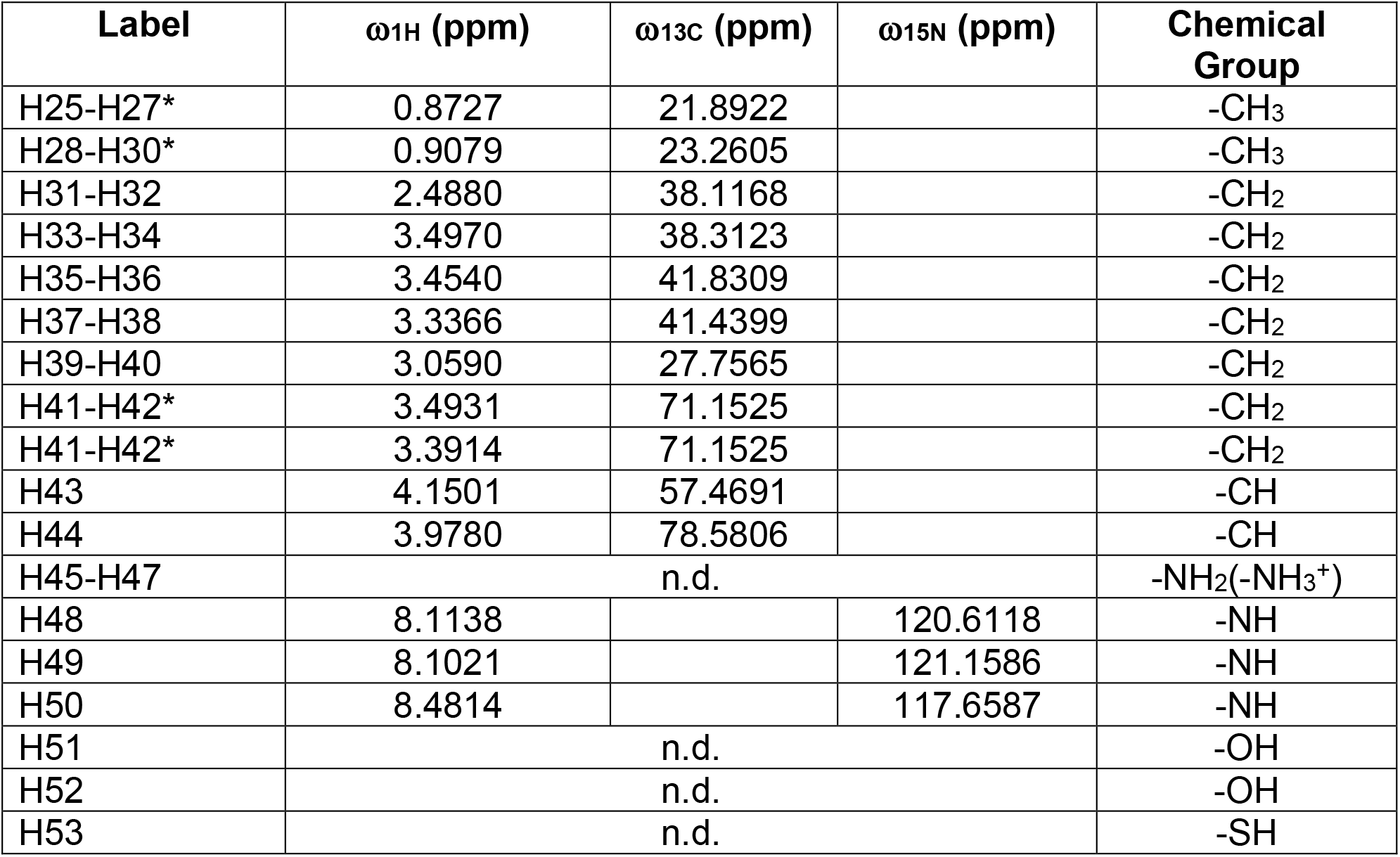
Chemical shift assignment of Cys-NH-pantetheine analog 1. Chemical shifts were referenced directly (^1^H) and indirectly (^13^C, ^15^N) with respect to DSS. n.d. = not detected. Atom labels marked with * indicate resonances that could not be uniquely assigned but were non-degenerate.

| Label | $\omega_{1\text{H}}$ (ppm) | $\omega_{13\text{C}}$ (ppm) | $\omega_{15\text{N}}$ (ppm) | Chemical Group |
| --- | --- | --- | --- | --- |
| H25-H27* | 0.8727 | 21.8922 |  | -CH<sub>3</sub> |
| H28-H30* | 0.9079 | 23.2605 |  | -CH<sub>3</sub> |
| H31-H32 | 2.4880 | 38.1168 |  | -CH<sub>2</sub> |
| H33-H34 | 3.4970 | 38.3123 |  | -CH<sub>2</sub> |
| H35-H36 | 3.4540 | 41.8309 |  | -CH<sub>2</sub> |
| H37-H38 | 3.3366 | 41.4399 |  | -CH<sub>2</sub> |
| H39-H40 | 3.0590 | 27.7565 |  | -CH<sub>2</sub> |
| H41-H42* | 3.4931 | 71.1525 |  | -CH<sub>2</sub> |
| H41-H42* | 3.3914 | 71.1525 |  | -CH<sub>2</sub> |
| H43 | 4.1501 | 57.4691 |  | -CH |
| H44 | 3.9780 | 78.5806 |  | -CH |
| H45-H47 | n.d. |  |  | -NH<sub>2</sub>(-NH<sub>3</sub><sup>+</sup>) |
| H48 | 8.1138 |  | 120.6118 | -NH |
| H49 | 8.1021 |  | 121.1586 | -NH |
| H50 | 8.4814 |  | 117.6587 | -NH |
| H51 | n.d. |  |  | -OH |
| H52 | n.d. |  |  | -OH |
| H53 | n.d. |  |  | -SH |

